# Most small cerebral cortical veins demonstrate significant flow pulsatility: a human phase contrast MRI study at 7T

**DOI:** 10.1101/2020.01.24.912329

**Authors:** Ian D Driver, Maarika Traat, Fabrizio Fasano, Richard G Wise

## Abstract

Phase contrast MRI has been used to investigate flow pulsatility in cerebral arteries, larger cerebral veins and the cerebrospinal fluid. Such measurements of intracranial pulsatility and compliance are beginning to inform understanding of the pathophysiology of conditions including normal pressure hydrocephalus, multiple sclerosis and dementias. We demonstrate the presence of flow pulsatility in small cerebral cortical veins, for the first time using phase contrast MRI at 7 Tesla, with the aim of improving our understanding of the haemodynamics of this little-studied vascular compartment. An automated method for establishing where venous flow is pulsatile is introduced, revealing significant pulsatility in 116 out of 146 veins, across 8 healthy participants, assessed in parietal and frontal regions. Distributions of pulsatility index and pulse waveform delay were characterized, indicating a small, but statistically significant (p<0.05), delay of 59±41 ms in cortical veins with respect to the superior sagittal sinus, but no differences between veins draining different arterial supply territories. Measurements of pulsatility in smaller cortical veins, a hitherto unstudied compartment closer to the capillary bed, could lead to a better understanding of intracranial compliance and cerebrovascular (patho)physiology.

## 1 Introduction

Pulsatility in cerebral veins is thought to be a passive process, a response to intracranial pressure changes arising due to arterial pulsatility through the cardiac cycle (Greitz et al., 1992). This normal process may be altered in some pathologies, such as normal pressure hydrocephalus, dementias, including Alzheimer’s disease, and multiple sclerosis (Bateman, 2000; Greitz, 2004; Mitchell, 2008; Henry-Feugeas and Koskas, 2012; Beggs, 2013; Bateman et al., 2016; Rivera-Rivera et al., 2017). Non-invasive measurements of venous pulsatility have the potential to provide insight into such pathologies. For example, reduced intracranial compliance has been observed in both normal pressure hydrocephalus and multiple sclerosis patients (Bateman et al., 2016). Intracranial compliance in this context refers to the capacity of the intracranial tissue to dissipate the arterial pulse wave, predominantly through cerebrospinal fluid (CSF) movements and the cerebral vasculature.

Phase contrast MRI (pcMRI) is a powerful tool for measuring blood velocity in cerebral blood vessels, and venous pulsatility has been observed previously (Bateman and Loiselle, 2007; Stoquart-Elsankari et al., 2009; Bateman et al., 2016; Rivera-Rivera et al., 2017) in studies limited to large veins, such as the venous sinuses and jugular veins and in large cortical veins, which drain directly into the sagittal sinus (Bateman, 2003). Applying these measurements to smaller cortical veins would allow changes to be studied in a venous compartment closer to the capillary bed and a better understanding to be gained of the functional consequences of venous pulsatility at the tissue level.

The primary objective of the work presented here is to assess whether pulsatility can be observed in small cortical veins. To this end, a cardiac gated 2D phase contrast MRI sequence (a standard vendor sequence) was acquired at 7 Tesla. Additionally, we introduce metrics to characterize this pulsatility in these veins, some of which are on the spatial scale of, or smaller than the in-plane image resolution. As such, we assess whether venous pulsatility can be resolved in small cortical veins by surveying over 100 cortical veins in 8 healthy participants and developing methods to characterize the pulsatility in these small blood vessels.

## 2 Material and Methods

Eight healthy participants (22-45 years; 4 female/4 male) took part in this study. The School of Psychology, Cardiff University Ethics Committee approved this study and subjects gave written informed consent prior to participating. This research was performed in accordance with the guidelines stated in the Cardiff University Research Integrity and Governance Code of Practice (version 2, 2018). Measurements were performed on a whole-body 7 Tesla research MR-system (Magnetom, Siemens Healthcare GmbH, Erlangen, Germany) with 32-channel head receive/volume transmit (Nova Medical, Wilmington MA). A whole-brain T_2_*-weighted 3D FLASH localizer image (0.6 mm isotropic, TR/TE = 18/11 ms; flip angle = 8°) was acquired for slice planning. A single pcMRI slice (0.6×0.6×5 mm; 192 mm FOV; TR/TE=11.55/6.94 ms; flip angle=10°) was positioned obliquely, approximately 2 cm above the corpus callosum and covering superior parietal and frontal regions (Figure 1a). The position was chosen to maximize the number of veins cutting transversally through the slice, with veins identified as hypointense in the localiser image. The pcMRI sequence was acquired with a 2D velocity encoding scheme, with v_enc_ = 10 cm/s in the through-slice direction. The value of 10 cm/s was chosen for sensitivity to slow-flowing small cortical veins (Figure 1b-c). An ungated (average over cardiac cycle) dataset was acquired first (24 s) to check slice positioning. A cardiac gated pcMRI dataset was acquired using prospective gating triggered using a photoplethysmogram with the sensor placed on the subject’s finger. The acquisition window was adjusted for each subject’s cardiac cycle duration and was set to be short enough to acquire all the cardiac phases (33-45 cardiac phases per subject, median=38) before the next heartbeat. Note that a single cardiac phase took two TRs to acquire (23.1 ms), consisting of a pair of flow-encoded and flow-compensated readouts. Three signal averages were acquired for each k-space line, giving an approximate acquisition time of 15 minutes. Additionally, 3D (0.6 mm isotropic, 60 slices) T_2_*-weighted FLASH (T_2_*w) and time of flight (TOF) images were acquired with the same centre and orientation as the pcMRI slice, for use in distinguishing veins from arteries. T_2_*w acquisition parameters: 192mm FOV; TR/TE=16/10ms; flip angle=8°; GRAPPA 2. TOF acquisition parameters: 192mm FOV; TR/TE=12/4ms; flip angle=20°; GRAPPA 2.

**Figure 1:**
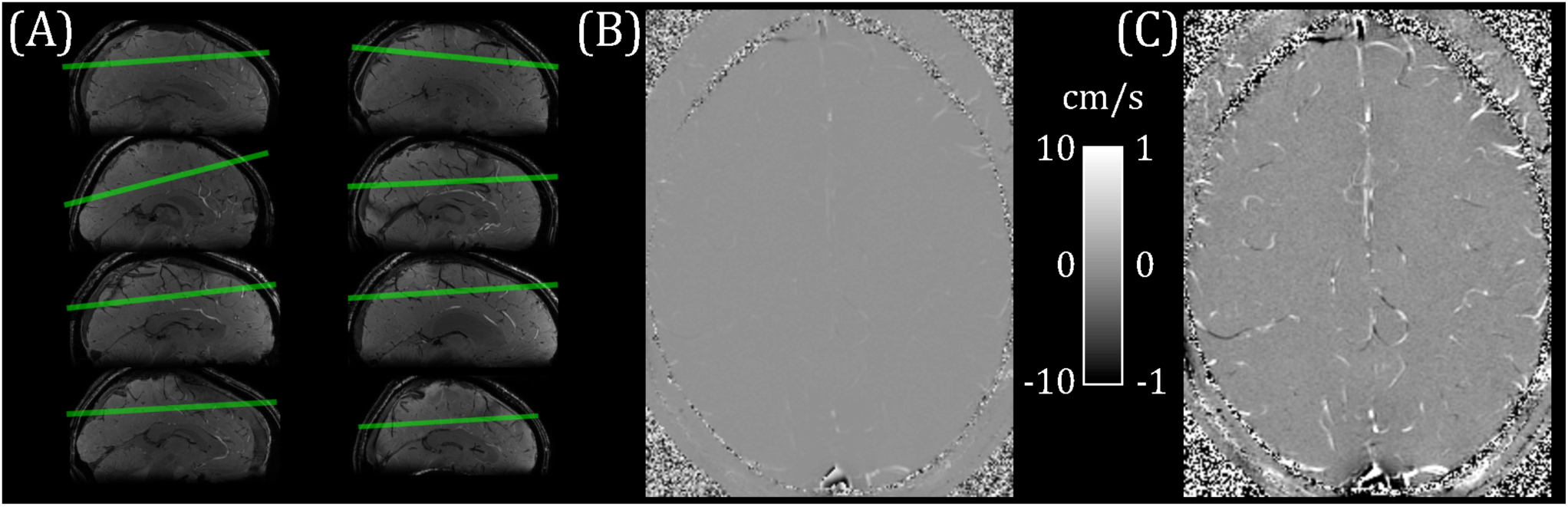
(A) pcMRI slice positioning for each subject (overlaid on a sagittal cut-through of the whole-brain T_2_*-weighted localizer image). (B) Example flow velocity map (mean across cardiac cycle). (C) Windowed version of (B), ranging from -1 cm/s to +1 cm/s.

Blood vessels were segmented using the pcMRI magnitude image (the magnitude of the complex difference between flow-compensated and flow-encoded images), averaged across the cardiac cycle. A threshold of two standard deviations above the image mean was applied to select small blood vessels. Individual vessels were indexed using 8-nearest-neighbour clustering, such that a super-threshold voxel is included in the same cluster as any of the 8-nearest-neighbour voxels that are also above the threshold (Figure 2a-c). Each cluster lying within the brain was manually characterized as a vein, an artery, or the superior sagittal sinus (Figure 2d-g). Veins and arteries were distinguished by overlaying these clusters onto the T_2_*w and TOF images, as follows. Veins appear hypointense (short T_2_*) and arteries appear hyperintense (inflow) on the T_2_*w image. The TOF image provides heightened inflow contrast, with arteries appearing bright and the majority of veins are invisible on the TOF image. Minimum intensity projections of T_2_*w (Figure 2e), maximum intensity projections of TOF (Figure 2g), both calculated over the extent of the 5mm pcMRI slice, and the original 0.6mm slice thickness T_2_*w (Figure 2f) and TOF images were used to identify whether the cluster represented a vein or an artery. Where this was unclear, or a vein and an artery both overlapped with the cluster, the cluster was discarded. Finally, veins were sub-divided into larger (3-5mm diameter) cortical veins, which appeared at the external cortical surface (hereafter referred to as surface veins), of the type reported previously (Bateman, 2003) and smaller cortical veins (∼0.6-2mm diameter), mostly lying at the intrasulcal cortical surface, which have not been studied previously (Figure 2h).

**Figure 2:**
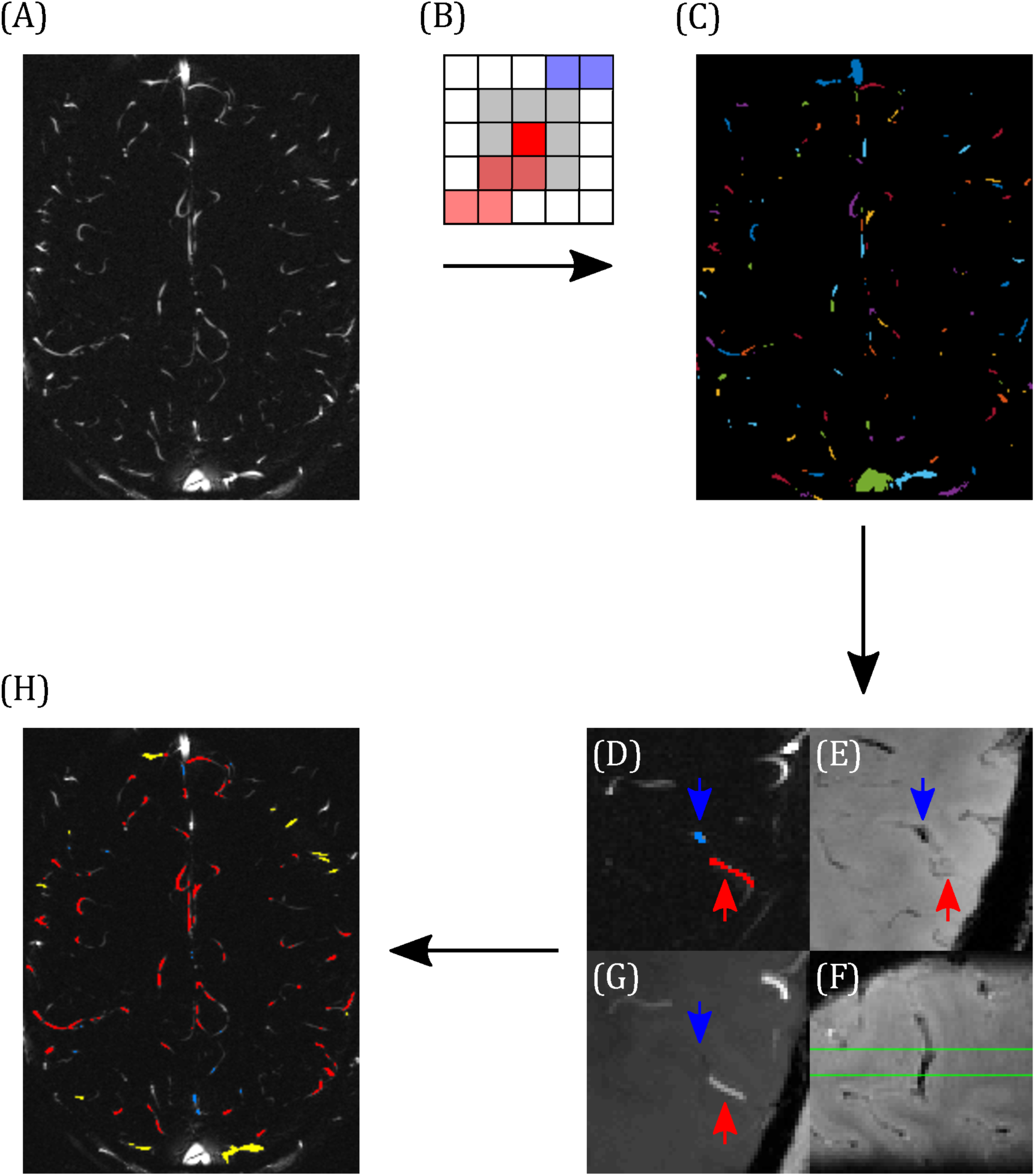
Demonstrating the classification of arteries and veins. (A) Example pcMRI magnitude image. (B) Clusters are formed such that each voxel with a pcMRI magnitude greater than two standard deviations above the mean is included in the same cluster as any of its 8 nearest neighbours (grey region) that are also above this threshold. In this example, red and blue voxels are separate clusters. (C) Example of the resulting cluster index map. (D-F) Example classification of a vein (blue) and an artery (red), with blue and red arrows used to identify the same locations across the images. (D) The two clusters of interest (blue and red), overlaid on the pcMRI magnitude image. (E) T_2_*w minimum intensity projection, showing a hypointensity at the blue cluster indicating a vein. (F) Sagittal slice of the T_2_*w image, centred on the blue cluster, showing the trajectory of the hypointense vein through the pcMRI slice (bound by the green lines). (G) TOF minimum intensity projection, showing a hyperintensity at the red cluster indicating an artery. (H) Example classification map overlaid on the pcMRI magnitude image. Arteries are shown in red and veins were sub-divided into large cortical veins, observed at the surface of the brain (yellow) and small cortical veins (blue).

For each cluster, flow velocity was calculated as follows from the pcMRI phase images (the phase difference between flow-compensated and flow-encoded images). A 10 mm (17×17 voxel) median filter was used to remove background phase offsets in the pcMRI phase images (Bouvy et al., 2016; Geurts et al., 2018). Voxelwise phase unwrapping was performed in the time (cardiac phase) dimension, such that phase jumps greater than π were unwrapped. Such wrapping only occurred in the superior sagittal sinus and arteries with high flow speeds. Taking the spatial mean across voxels for each cluster, then scaling by v_enc_/π to convert from radian units to cm/s calculated the flow velocity component through the slice.

The sagittal sinus pulse waveform was calculated from where the sagittal sinus cuts through the posterior portion of the slice. The top 20 voxels by intensity in the pcMRI magnitude image, averaged across the cardiac cycle, were selected. Median filtering and temporal unwrapping of the pcMRI phase dataset were performed, as described above. The high flow speed of the sagittal sinus meant that the signal phase in most of the voxels wrapped around multiple times. In this study, the sagittal sinus waveform was required for assessing timing differences, rather than absolute flow speeds, therefore unwrapping in space was performed to move the signal phase from all 20 voxels into the same wrapping point, rather than the true flow speed. The posterior portion of the sagittal sinus flowed superior to inferior, encoding a negative signal phase. Therefore, unwrapping in space was accomplished by subtracting 2π from any voxels with positive signal phase. Taking a median across the 20 voxels and scaling by v_enc_/π to convert from radian units to cm/s calculated the sagittal sinus pulse waveform.

Venous pulse waveforms were characterized firstly by detecting the presence of pulsatility based on a statistical criterion, then by calculating pulsatility index (PI) and temporal lag of the waveform relative to the sagittal sinus. In order to assess whether a waveform was pulsatile, we define a statistical parameter, pulsatility contrast to noise ratio (PCNR), as follows. It was assumed that lower frequency cardiac cycle-locked variations in the flow waveform were pulsatility, whereas higher frequency variations were noise. Waveforms were low-pass filtered (Savitzky-Golay; 3^rd^ order, 15 timepoint frame size). The velocity range (*Δv*) was taken as the range of this filtered waveform. PCNR was defined as the ratio of *Δv* and the standard deviation of the residuals between unfiltered and filtered timepoints (*res*) – see equation 1 and Figure 3:

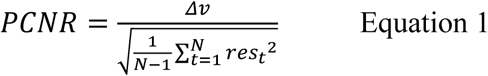

**Figure 3:**
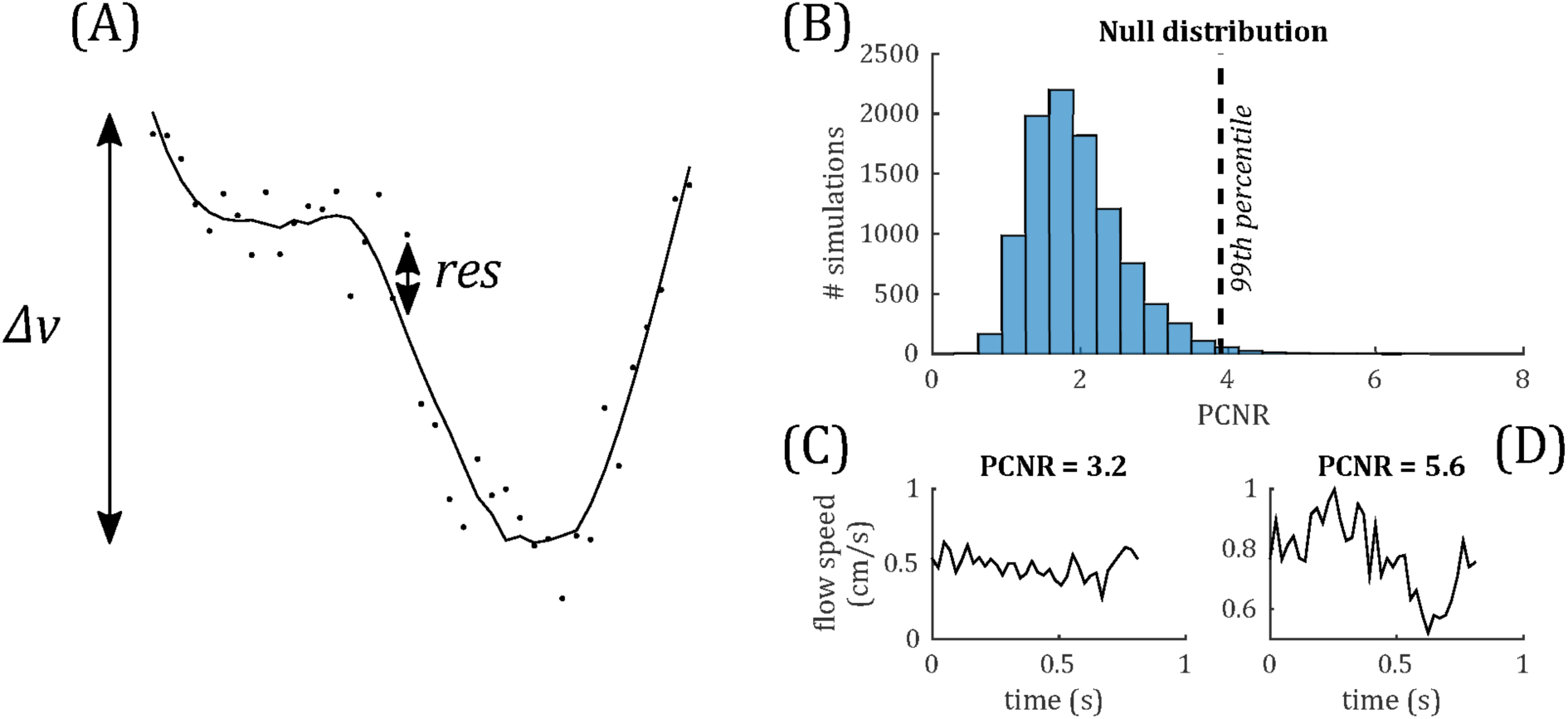
(a) Demonstrating definition of the PCNR parameter (see equation 1), with filtered curve (line) overlaid on original waveform (dots). PCNR is the ratio of the range of the curve *Δv* and the standard deviation of the residual (*res*). (b) Monte Carlo simulations, calculating PCNR from Gaussian noise waveforms. The 99^th^ percentile corresponding to PCNR > 3.9, whilst the distribution remains unaffected by changes in noise level or number of timepoints (data not shown). (c) example waveform which is not classed as pulsatile (PCNR ≤ 3.9) and (d) one that is (PCNR > 3.9).

PCNR is analogous to a t-statistic and, unlike PI, PCNR is not biased by low mean values. A statistical criterion for pulsatility was set as PCNR > 3.9 based on Monte Carlo simulations of the PCNR null distribution (see Figure 3), such that a threshold of PCNR > 3.9 corresponds to p<0.01. PI was calculated as *Δv* / mean(|*v*|).

Temporal lag was calculated relative to the superior sagittal sinus by calculating the peak cross-correlation between the unfiltered vein and sagittal sinus flow velocity waveforms, as demonstrated in Figure 4. For comparison, the temporal lags and PCNR of artery cluster waveforms were also calculated by the same method as for veins. PI was not assessed for arteries because the 10 cm/s v_enc_ was optimized for the flow speed of small veins. Higher arterial flow speeds caused signal phase wrapping, resulting in inaccurate mean flow velocity and PI estimates for arteries.

**Figure 4:**
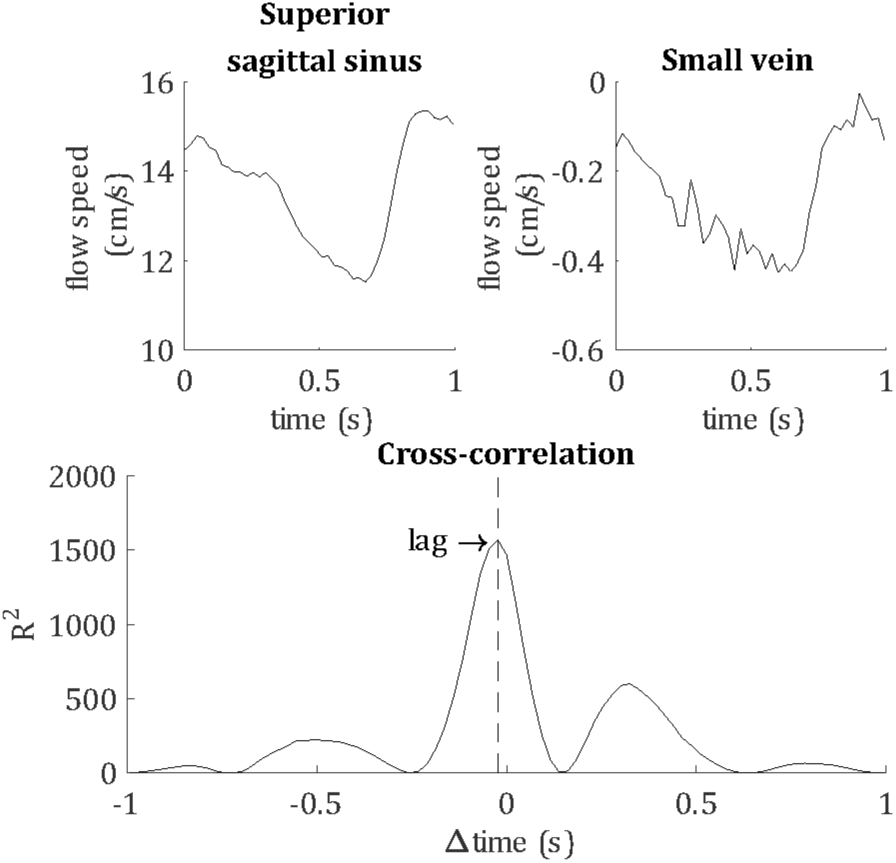
Demonstrating the calculation of the temporal lag of each blood vessel, relative to the superior sagittal sinus. For this example, flow velocity waveforms are shown for the superior sagittal sinus and a small vein from a single subject. The lag of the pulse waveform of each blood vessel was calculated as the peak cross-correlation with the sagittal sinus waveform, as shown by the vertical dashed line.

To explore spatial variations in the venous pulsatility characteristics, vessels were considered according to the local feeding arterial territory in which they lay, i.e. anterior, left-middle, right-middle or posterior cerebral artery. Feeding arteries from the four vascular territories were clearly separable in the TOF image and four regions were manually drawn at the level of the pcMRI slice, making reference to literature example arterial territory maps such as those found in Mut et al. (2014) and Hart et al. (2018). Vessel clusters lying within each region were labeled accordingly.

## 3 Results

146 veins were identified across the 8 subjects (range 12-26 veins per subject), of which 116 showed pulsatility (range 8-21 veins per subject), determined by PCNR > 3.9. Subject-averaged venous flow velocity waveforms are presented in Figure 5 for all veins that show pulsatility. The equivalent plot for the remaining veins with PCNR ≤ 3.9 is presented in Supplementary Figure S1. When these ‘non-pulsatile’ veins are averaged together within each subject, pulsatility appears present in 4 of the subjects. This suggests that at least some of these ‘non-pulsatile’ veins are pulsatile and could be resolved with a higher signal-to-noise ratio. For comparison, out of the 432 arteries identified, 385 showed pulsatility. Sub-dividing the veins, 59 out of 69 identified surface veins (3-5 mm diameter) and 57 out of 77 smaller cortical veins (<2 mm diameter) were pulsatile. Qualitatively, there was no clear trend in the orientation, tortuosity or location of the non-pulsatile veins.

**Figure 5:**
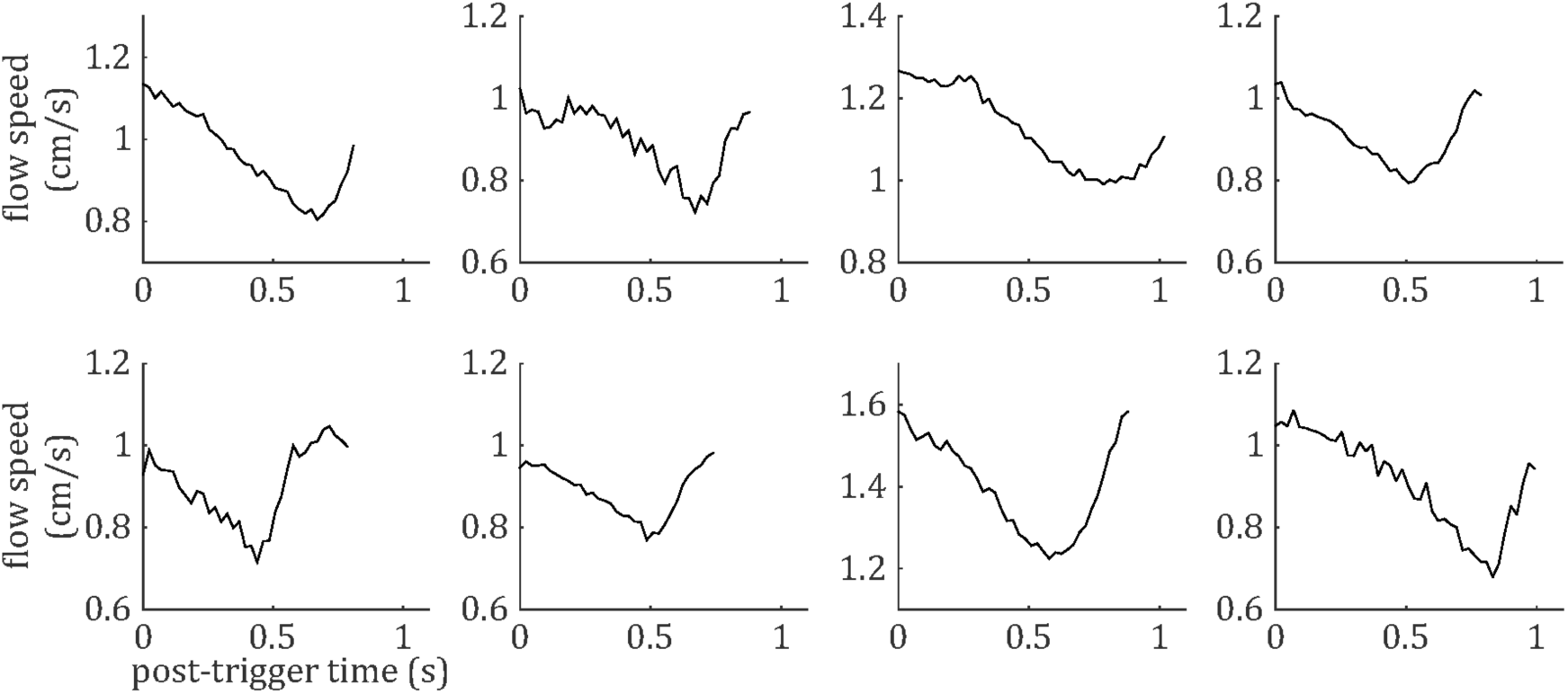
Cardiac cycle synchronized venous blood flow time-courses for each subject (mean across all pulsatile cortical veins; N.B. does not include the venous sinuses).

The distributions of PCNR, PI and temporal lag across all veins from all subjects are presented in Figure 6, split into smaller and surface veins. The distributions appear skewed, so summary values of PI and temporal lag were calculated for each subject by taking the median across vessels. These summary values are reported as follows: PI = 0.30±0.05 (mean±standard deviation across subjects) when all veins were combined. PI values sub-divided into small cortical veins and larger surface veins are presented in Figure 7. A comparison of PI between small cortical veins and larger surface veins did not reach significance (t(7) = 1.56; p = 0.16; 2-tailed paired t-test across participants). For all veins combined temporal lag = 59±41 ms and for arteries temporal lag = 23±105 ms, both with respect to the superior sagittal sinus. Temporal lag values sub-divided into small cortical veins and larger surface veins and for arteries are presented in Figure 7. Small cortical veins (t(7) = 3.43; p_corr_ = 0.044) and all veins combined (t(7) = 4.11; p_corr_ = 0.018) showed a significant delay with respect to the superior sagittal sinus, when temporal lag was compared to zero with a 2-tailed t-test and Bonferroni corrected for 4 comparisons (small veins, surface veins, all veins combined and arteries). For completeness, the temporal lags of surface veins (t(7) = 3.21; p_corr_ = 0.059) and arteries (t(7) = 0.62; p_corr_ > 1) were not significantly different from the superior sagittal sinus. A comparison of temporal lags between smaller cortical veins, larger surface veins and arteries did not reach significance (F(2,21) = 0.57; p = 0.58).

**Figure 6:**
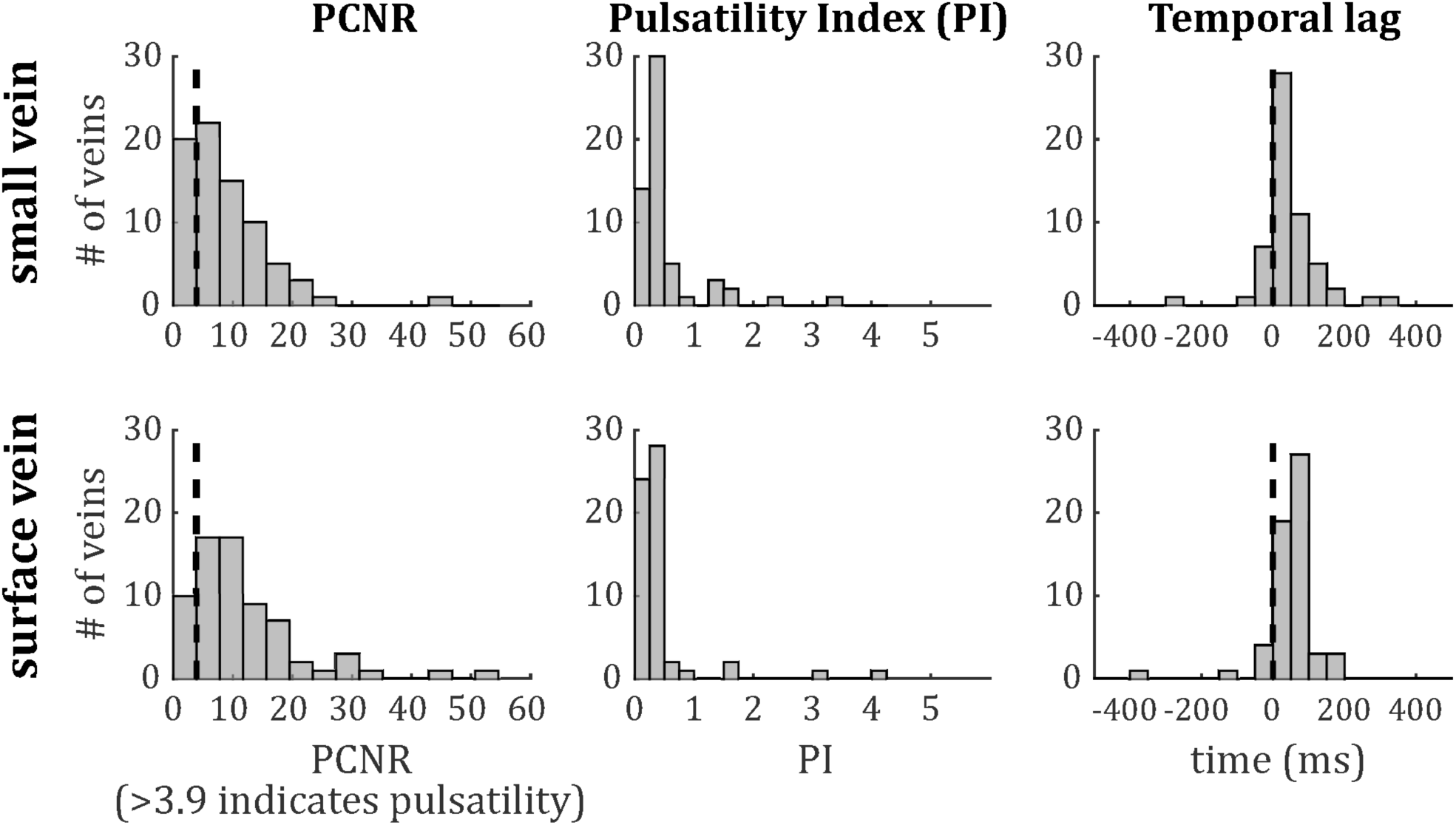
Histograms of (left) PCNR, (middle) PI and (right) temporal lag for small veins (top) and surface veins (bottom). For PCNR, all veins are included, whereas for PI and temporal lag, only pulsatile veins (PCNR >3.9) are included. The dashed line in the left panels corresponds to the PCNR = 3.9 threshold. The dashed line in the right panels indicates zero lag compared to the superior sagittal sinus.

**Figure 7:**
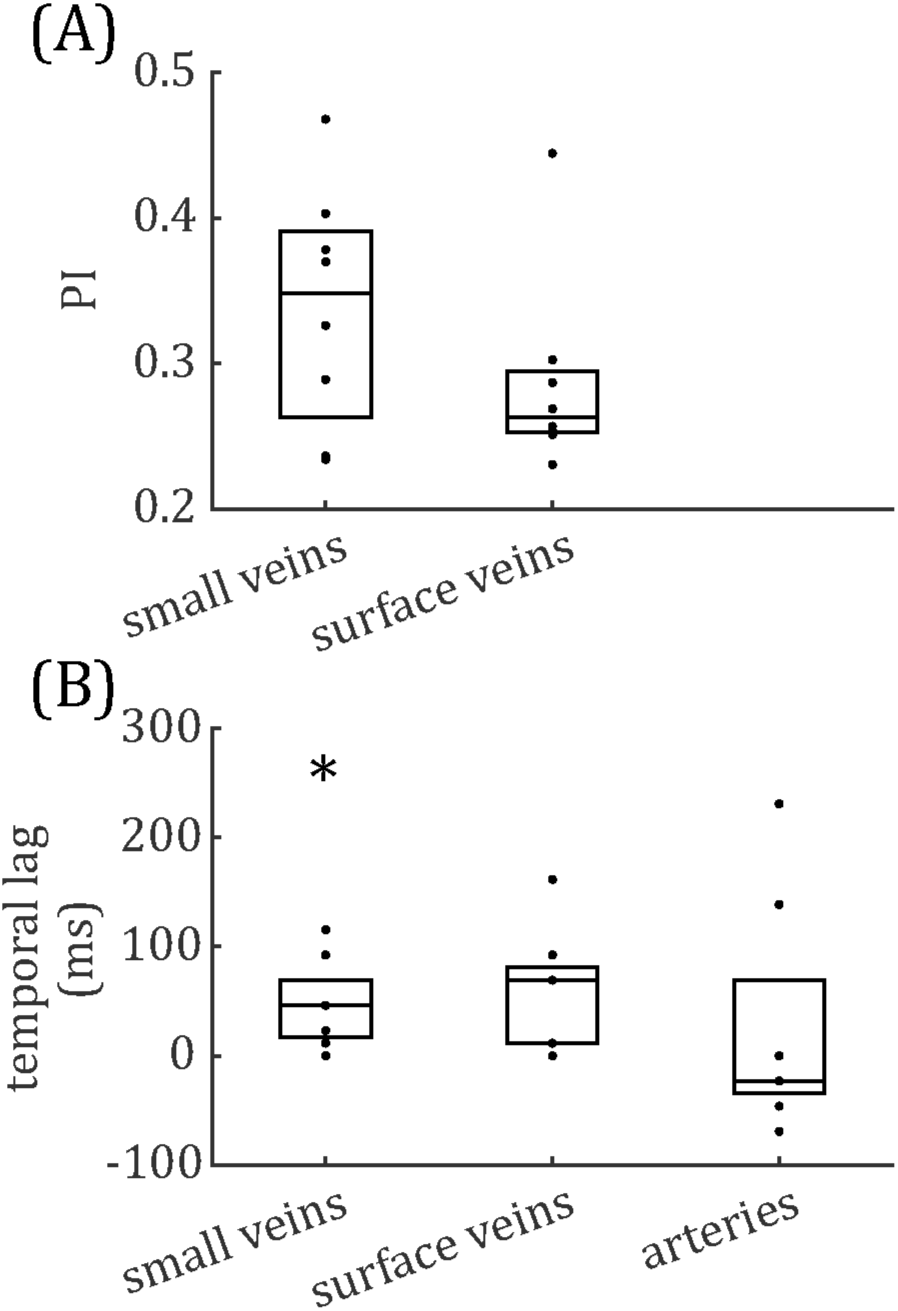
Boxplots for (A) pulsatility index (PI) and (B) temporal lag, relative to the superior sagittal sinus. Boxes show median and interquartile range across subjects and each subject’s data (median across vessels) is presented as a dot. * Bonferroni-corrected p <0.05 that the temporal lag is different from zero.

Temporal lag compared across the four vascular territories (median across vessels within each subject, one-way ANOVA across subjects) did not show significant inter-regional differences for either veins (F(3,27) = 0.39; p = 0.76) or arteries (F(3,28) = 0.13; p = 0.94). Vein PI compared across the four vascular territories did not reach significance (F(3,27) = 0.68; p = 0.57). Note that no pulsatile veins were identified in one territory (anterior cerebral artery) for a single subject, so this was treated as missing data in each ANOVA. Histograms of temporal lag for all veins and all arteries across all subjects are presented split into vascular territories in Figure 8.

**Figure 8:**
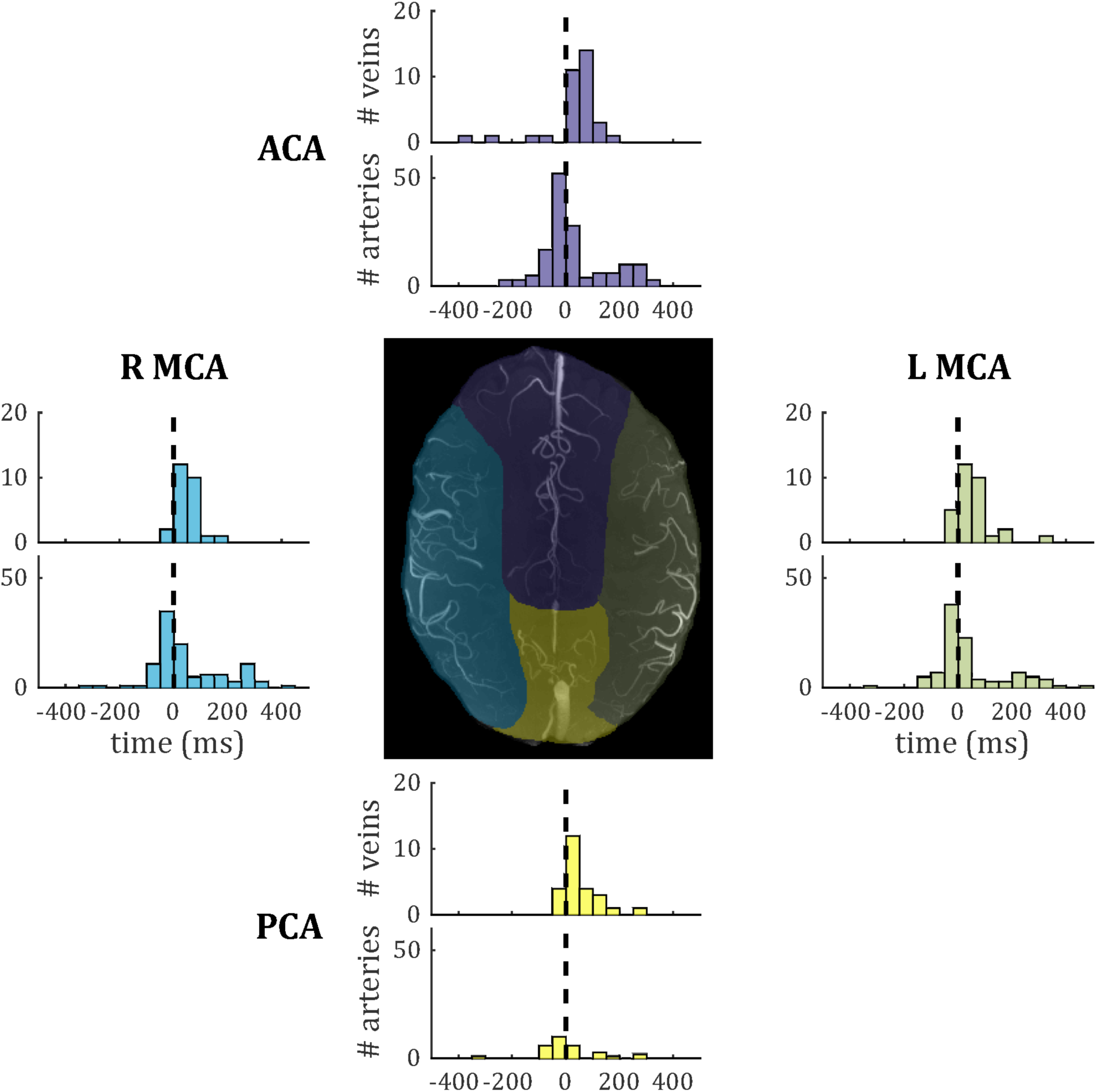
Histograms of temporal lag for all pulsatile veins (PCNR > 3.9) and all pulsatile arteries (PCNR > 3.9) within each vascular territory. For reference, the dashed line marks zero lag with respect to the superior sagittal sinus. Vascular territories are characterised by the regions broadly supplied by the anterior cerebral artery (ACA), left- and right- middle cerebral artery (L MCA; R MCA) and posterior cerebral artery (PCA). An example territory map overlaid on TOF maximum intensity projection is shown at the centre, with colour of each territory matching the histograms for vessels from that territory.

## 4 Discussion

We have directly observed pulsatility in small cortical veins using pcMRI at 7T with v_enc_= 10 cm/s. Although a similar observation has been made previously in similar-sized cerebral arteries (Bouvy et al., 2016; Schnerr et al., 2017; Geurts et al., 2018), we believe this to be the first reported observation of pulsatility in small cortical veins, upstream of the large cortical veins draining directly into the superior sagittal sinus reported previously (Bateman, 2003). We extend this novel finding by surveying 146 veins across 8 subjects and develop methods to characterize pulsatility in these blood vessels, which are on the order of the 0.6 mm in-plane voxel size. 116 out of these 146 veins (79%) were characterized as pulsatile. Finally, we survey the distribution of PI and temporal lag across this healthy population of cortical veins.

This work establishes that small cortical veins are pulsatile, however, it does not establish how these measurements vary across a healthy population, or to what extent this pulsatility can be perturbed, either with physiological stimulus or pathophysiology. The observation of pulsatility is robust across the large number of veins studied, observed in 116 of the 146 veins studied and seen in multiple veins in all 8 subjects studied. Therefore the data presented here introduces the phenomenon of pulsatile small cortical veins and future studies are required to establish test-retest reproducibility and perturbations of the phenomenon.

Our measurement of small vein pulsatility can be applied to provide new information for understanding the role of veins in pathologies with altered intracranial haemodynamics and compliance. Normal pressure hydrocephalus is associated with decreased intracranial compliance (Bateman, 2000; Bateman and Loiselle, 2007; Wagshul et al., 2011; Beggs, 2013; Bateman et al., 2016), which manifests in the sagittal sinus pulse waveform as a decrease in arteriovenous delay (AVD). Similar evidence for decreased intracranial compliance has been observed in multiple sclerosis (Bateman et al., 2016) and Alzheimer’s Disease (Rivera-Rivera et al., 2017). Proposed pathophysiological mechanisms for this decreased intracranial compliance include impaired CSF outflow or absorption (Greitz, 2004), venous hypertension (Bateman, 2008; Wagshul et al., 2011; Beggs, 2013), or breakdown of the Windkessel mechanism leading to the pulse wave propagating through the capillary bed (Bateman, 2008; Beggs, 2013). Our measurement of small vein pulsatility can be used to study these pathophysiological mechanisms closer to the level of the capillary bed, whereas the venous sinuses and jugular veins are further downstream.

The results presented here have the potential for informing the future study of venous function, intracranial pulsatility and compliance (Wagshul et al., 2011). These parameters can be measured using direct intracranial pressure measurements of pressure pulsatility, or using transcranial Doppler ultrasound or pcMRI to measure flow pulsatility. Short of invasive intracranial pressure measurements, intracranial compliance is typically inferred either through measurements of arterial and venous flow-velocity pulse waveforms, using the AVD as a surrogate measure of compliance (Bateman et al., 2016), or through comparing arterial and CSF pulse waveforms (Baledent et al., 2001) and fitting a transfer function to calculate intracranial pressure (Alperin et al., 2000). The AVD method is typically calculated based on measurements in large blood vessels feeding and draining the brain, such as the carotid arteries and venous sinuses. Therefore it is sensitive to any changes in arterial, venous and microvascular compliance. The method using measurements of CSF flow velocity relies on measuring low flow velocities in CSF, with the associated lower signal-to-noise ratio. The small veins that are the focus of this study are closer to the tissue of interest than the venous sinuses or cerebral aqueduct.

To place our results in context with the literature, venous flow speeds observed of the order 1 cm/s are within the range predicted previously (Piechnik et al., 2008), and flanked by flow velocity values reported in basal ganglia and centrum semiovale arteries (Bouvy et al., 2016; Geurts et al., 2018). The pulsatile waveforms of the veins observed here have a similar shape to those reported in basal ganglia and centrum semiovale arteries (Bouvy et al., 2016). Cortical vein PI = 0.30±0.05 is within the range of 0.2-0.5 reported previously in the superior sagittal sinus (Stoquart-Elsankari et al., 2009; Rivera-Rivera et al., 2017) and the PI values of 0.40±0.09 and 0.28±0.07 reported previously in basal ganglia and centrum semiovale arteries, respectively (Bouvy et al., 2016). To put these PI values into perspective, internal carotid artery PI of 0.8-0.9 has been reported previously (Rivera-Rivera et al., 2017; Bouillot et al., 2018).

Pulsatility was characterized in 146 veins, allowing an initial survey of PI and temporal lag to be performed. Both showed non-Gaussian distributions, with positive skews (Figure 6). To further assess these distributions, comparisons were made in PI and temporal lag by sub-dividing veins into small cortical veins and larger surface veins, and in temporal lag by comparing veins to arteries. Vessels were also sub-divided by vascular territory. To avoid confounding comparisons by mixing intra- and inter-subject variability, summary statistics of median across veins for each subject were used for these comparisons, which did not reach significance. The small number of subjects combined with inter-subject variability (Figure 7) limit the power of these comparisons to detect small differences. However, Figures 7 and 8 do suggest hints of small veins having higher PI than surface veins and of an earlier arterial and later venous temporal lag (with their modes lying either side of zero in Figure 8). A limitation of the temporal lag measurement is the sensitivity of the cross-correlation to noise in the superior sagittal sinus waveform, which has potential to introduce subject-specific bias in the temporal lag reported. This could explain the large variance in artery temporal lags observed in this study (Figures 7b and 8). For example, the second histogram peak at ∼ 200 ms in arterial temporal lag is largely due to a single subject, which could be driven by a noisy sagittal sinus waveform.

An interesting finding is that the flow waveform from the superior sagittal sinus leads the flow waveform of small cortical veins, despite being downstream of these veins. A possible explanation for this is that the velocity wave-front propagates back upstream from the superior sagittal sinus towards the smaller veins sampled. This observation, along with the small arterial waveform leading that of the superior sagittal sinus, would be consistent with the effect of the arterial pressure wave, and associated increase in arterial volume, resulting in a decrease in superior sagittal sinus volume first, followed by the smaller veins. Such a mechanism implies transmission of the arterial pressure pulse through the brain tissue. An alternative explanation of direct transmission of the arterial pressure pulse through the arteries, through the capillaries and on to the veins (Hahn et al., 1996; Rashid et al., 2012) is perhaps less likely given the observations.

Technical considerations of this study include partial volume errors, arising due to the focus on small veins whose diameter is on the order of the in-plane voxel size and considerably smaller than the slice thickness, resulting in veins occupying only a fraction of the voxel volume. This produces a large uncertainty in the cross-sectional area, which would be required for measurements of absolute flow and stroke volume in the veins. Furthermore, most vessels will not pass perpendicularly through the image slice, and thus not parallel to the flow encoding direction, reducing the detectability of motion. These partial volume considerations are likely to reduce the detection of pulsatility rather than to result in false positives. In addition, compliance-based changes in this voxel volume fraction across the cardiac cycle will result in changes in the signals measured that are not directly related to flow speed. This could bias PI measurements. Changes in cross-sectional area across the cardiac cycle have been observed in the middle cerebral artery (Warnert et al., 2016), but to best of our knowledge have not been reported in the small cortical veins and arteries studied here. Cortical veins may be expected to show a smaller change in cross sectional area than cortical arteries (Lee et al., 2001; Kim et al., 2007; Chen and Pike, 2009, 2010; Wesolowski et al., 2019).

Use of 10 cm/s *v*_*enc*_ maximized dynamic range for measuring flow speed in the slow-flowing cortical veins at the expense of being able to quantify flow in blood vessels supporting faster flow, such as arteries and the sagittal sinus. Future studies interested in all of these blood vessels will either need to include multiple acquisitions with different *v*_*enc*_ values optimized for each type of blood vessel, or work to find an intermediate *v*_*enc*_, sacrificing contrast for slow flowing veins to ensure the signal does not wrap in faster flowing blood vessels. The manual identification of veins and arteries is subject to inter-observer differences and is labour intensive. The small number of subjects studied here means that inter-subject variability cannot be reliably assessed in this study. Cerebral blood pressure is modulated by factors such as age, gender, hydration level and caffeine intake, so these would also need to be controlled for when investigating inter-subject variability.

This study acquired three signal averages for each k-space line, making acquisition time approximately 15 minutes. However, the signal-to-noise ratio was more than sufficient to clearly resolve pulsatility in small cortical veins, so a single average could be used in future, reducing the acquisition time to 5 minutes. Employing acceleration techniques, such as parallel imaging and view sharing, could further shorten the acquisition time. This study uses a 2D acquisition and a single v_enc_ direction, which limits the coverage of small veins, so future work will assess undersampled 3D pcMRI approaches (Rivera-Rivera et al., 2017).

In conclusion, we report the first observation of pulsatility in small (∼0.6-2mm) cortical veins in-vivo in humans. We present methods to characterize pulsatility in small blood vessels and present a survey of the distributions of pulsatility in cortical veins in the healthy brain. This measurement has implications for studies of cerebral venous function and offers the potential for more detailed investigation of cerebral haemodynamics, cerebral compliance and intracranial pressure.

## 5 Abbreviations

AVD: Arteriovenous delay
PCNR: Pulsatility contrast to noise ratio
CSF: Cerebrospinal fluid
pcMRI: Phase contrast
MRI PI: Pulsatility index
TOF: Time of flight angiography
v_enc_: Velocity encoding gradient cutoff velocity

## 6 Acknowledgements

ID and RW acknowledge the generous support of the Wellcome Strategic Award, “Multi-scale and multi-modal assessment of coupling in the healthy and diseased brain,” grant reference 104943/Z/14/Z. The UK Medical Research Council (MR/M008932/1), the Welsh Government and Cardiff University provide funding support for the CUBRIC ultra-high field imaging facility.

## 7 Author Contribution Statement

ID and RW conceived and designed the research. ID, MT and FF acquired the data. ID and MT performed data analysis. ID and RW interpreted the results. ID wrote the manuscript and all authors provided critical revisions.

