## Supplementary Figure S1 for "Most small cerebral cortical veins demonstrate significant flow pulsatility: a human phase contrast MRI study at 7T"

### *Supplementary Material*

#### 1 Supplementary Figures

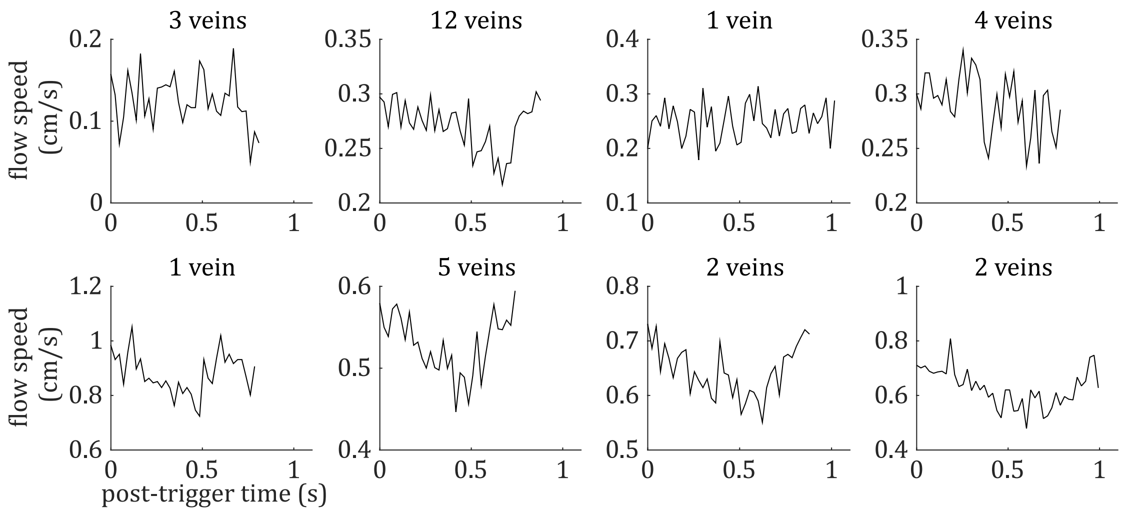


**Supplementary Figure 1.** Cardiac cycle synchronized venous blood flow time-courses for each subject for veins that do not meet the PCNR>3.9 threshold (i.e. 'non-pulsatile veins'). The number of non-pulsatile veins for each subject is displayed above the respective plot.
